# Comparison of environmental DNA and bulk DNA metabarcoding for assessing terrestrial arthropod diversity across three habitat types on Guam

**DOI:** 10.64898/2026.02.02.703366

**Authors:** Pritam Banerjee, Samantha Al-Bayer, Jerilyn Calaor, Sven Weber, Natalie Graham, Jeremy C. Andersen, Evan P. Economo, Susan Kennedy, Henrik Krehenwinkel, Rosemary Gillespie, George Roderick, Haldre Rogers, Kenneth P. Puliafico

**Author notes:** These authors contributed equally and jointly supervised this work (Haldre Rogers, Kenneth P. Puliafico).

## Abstract

DNA based methods offer a rapid and cost-effective way for detecting species occurrence and monitoring biodiversity; among them bulk DNA metabarcoding is well-established, and recently developed environmental DNA (eDNA)-based methods offer a non-destructive alternative. With a goal to develop suitable methods for assessing insect biodiversity in ecosystems for which DNA reference libraries are not well developed and incomplete, such as remote islands, we compared established bulk DNA metabarcoding methods with eDNA across three replicated terrestrial ecosystem types (limestone forest, degraded forest, and grassland) in Guam. Using two mitochondrial COI primer pairs, we performed bulk DNA metabarcoding of standard entomological collection methods (malaise traps, pan traps, vegetation beating), and compared the assessment of biodiversity with that from different eDNA sources (flowers, spider webs, leaves, tree trunks). In our samples, eDNA and bulk DNA metabarcoding both detected a large proportion of overall taxa (OTUs, 86.6% and 60.3%, respectively). Although DNA metabarcoding detected significantly more taxa, eDNA proved to be a reasonable non-destructive alternative. As expected, because of limitations in existing reference databases for remote habitats, species-level identification was achieved for only a few OTUs. Overall, the sampling approach was the dominant driver of arthropod diversity, explaining ∼17% of the observed variation, while habitat type accounted for ∼4%. Thus, each sampling approach captured some unique diversity signals and contributed to the complementary effect of maximizing detection. For rapid insect biodiversity surveys of terrestrial arthropods, we recommend an integrated metabarcoding approach, and in sensitive habitats where insect capture is undesirable, eDNA offers a powerful alternative to monitor diversity and community change.

## Introduction

Biodiversity is facing threats worldwide (Pereira et al., 2024), and we continue to lack sufficient expertise to implement fast and effective monitoring of known species (Baird & Hajibabaei, 2012), let alone undescribed taxa (Meier et al., 2025). Arthropods are a particularly understudied group and, notwithstanding concerning reports on their decline (Basset & Lamarre, 2019), we have limited data to infer their current threat status for many regions of the world (Sánchez-Bayo & Wyckhuys, 2019; Wagner, 2020).

Traditional entomological surveys use a variety of collecting approaches to capture whole organisms, which are then identified by experts. The collecting approaches include (i) malaise traps – non-attractant and static traps, primarily used for collecting flying insects, particularly those that are active during the day as well as those flying at night (Skvarla et al., 2021; Seymour et al., 2024; Buchner et al., 2025); (ii) beating vegetation that targets plant-associated arthropods by shaking or beating plants and capturing falling specimens (Kennedy et al., 2023); (iii) pan traps (white and yellow pan) that consist of small, brightly colored bowls filled with soapy water to passively attract and capture insects, particularly those that are visually attracted to specific colors (Westerberg et al., 2021). While these methods are effective in capturing a wide range of species, identifying insects morphologically following collection is time-consuming, laborious, and fails to keep up the speed for monitoring insects with rapid environmental changes and decline (Chua et al., 2023). Furthermore, the need for specialized taxonomic skills for each insect group makes the process even slower (Kim & Byrne, 2006). Moreover, given the large proportion of undescribed species (Berenbaum, 2017), the task is simply not impossible for many regions of the world.

As an alternative to traditional approaches, biomonitoring with DNA metabarcoding of the bulk samples (e.g., bulk DNA extraction from malaise traps, pan traps, and vegetation beating) is effective for rapidly detecting spatio-temporal patterns of change in insect diversity (Graham et al., 2023; Strutzenberger et al., 2024), and requiring limited taxonomic expertise (Yu et al., 2012). These bulk DNA metabarcoding techniques are now well-established in insect monitoring (Piper et al., 2019; Liu et al., 2020; Chua et al., 2023) and have been demonstrated to deliver satisfactory results (Watts et al., 2019). Additionally, recent development of environmental DNA (eDNA) based metabarcoding methods allows us to capture a wide range of taxa from environmental samples without collecting whole organisms, thus making species monitoring even faster and more cost-effective (Thomsen & Willerslev, 2015; Deiner et al., 2017). The eDNA-based method is widely used in aquatic environments, but its application in terrestrial environments is relatively new (Thomsen & Sigsgaard, 2019; Valentin et al., 2020; Pumkaeo et al., 2021; Gregorič et al., 2022), and is suitable for monitoring species under conservation management and/or in protected areas, where non-destructive monitoring is required (Thomsen & Willerslev, 2015; Beng & Corlett, 2020).

Here, we aim to understand the effectiveness of eDNA and bulk DNA metabarcoding in assessing arthropod diversity on Pacific islands, which are generally more susceptible to the rapid decline of species compared to most continental habitats (Fordham & Brook, 2010). We focus specifically on Guam, a small island (213 km²) at the southern end of the Mariana Island arc, which has been heavily impacted by invasive species and the gradual decline of native species (Fritts & Rodda, 1998). This combination of endemism, intense invasion pressure, and limited baseline data makes Guam an ideal model system for evaluating biodiversity monitoring approaches under real-world conservation constraints (Gressitt, 1994; Fritts & Rodda, 1998).

Many islands and some fragmented mainland systems face similar conditions, including rapid species turnover, strong biosecurity concerns, and ethical or logistical limitations on destructive sampling. As such, methodological insights gained from Guam are broadly transferable to other Pacific islands and other tropical environments where rapid, scalable, and minimally invasive monitoring tools are urgently needed. Improving monitoring strategies that can deliver timely, comparable data is critical for enabling early detection, adaptive management, and conservation action in these systems.

For such habitats, the question remains of how well newly developed sources of eDNA perform compared to bulk DNA metabarcoding methods in assessing overall taxonomic diversity? To answer this question, we designed a comprehensive study where eDNA methods were selected to match bulk DNA metabarcoding methods, and then compared, across habitat types (intact limestone forest, degraded forest, grassland) on Guam that differ in terms of anthropogenic disturbance (Figure 1, Table 1), with the goal of evaluating the utility of the approach in revealing differences between sites. We selected eDNA sources that might be expected to largely overlap with bulk collection approaches. For example, (i) eDNA recovered from flowers can capture a variety of flower visitors (Thomsen & Sigsgaard, 2019), which potentially overlaps with the insects caught in pan and malaise traps; (ii) eDNA extraction directly from spider webs can act as a natural air eDNA sampler (Gregorič et al., 2022), and could potentially capture a wide variety of airborne eDNA, which also can overlap with pan and malaise traps. Additionally, (iii) eDNA from surface washing of plant leaves and (iv) trunk (bark) can capture a variety of foliage-associated and/or canopy-dwelling arthropods interacting with plants (Valentin et al., 2020), which potentially overlaps with arthropods captured through vegetation beating.

**Table 1.** Sampling methods used and target arthropods in comparison of bulk DNA and eDNA metabarcoding.

| Traditional DNA-Metabarcoding sampling methods | eDNA Metabarcoding methods | Target arthropods |
| --- | --- | --- |
| Malaise traps | Flower sampling and Spider webs | Flying and crawling arthropods |
| Beating vegetation | Leaf wash water | Mobile, plant-dwelling arthropods |
|  | Trunk/surface rolling |  |
| Pan traps (yellow & white) | Flower sampling and spider webs | Pollinators |

## Material and methods

### Study Sites

Samples were collected in March and April 2023 from a total of nine sites representing three habitat types across the U.S territory of Guam (Figure 1). Two of the habitats were different forest types and classified as (i) intact limestone forest (sites A3, M3, M4), characterized by karst with fissures, caves, and jagged terrain and dominant plants, including *Ficus prolixa, Ochrosia oppositifolia, Pandanus tectorius, P. dubius,* and *Aglaia mariannensis*, and (ii) degraded forest (sites A1, A2, N1, M1), classified as mixed introduced forests, characterized by formerly cleared or developed lands left to regenerate naturally or planted, and now consisting of a mix of non-native and native plants. Common plants found in the mixed introduced forests include the non-native species *Leucaena leucocephala*, *Vitex pariviflora*, and *Acacia auriculiformis,* along with the native species *Meiogyne cylindrocarpa, Aglaia mariannensis,* and both *Pandanus* species. The third habitat was (iii) grassland (sites N2, M2), classified also as badlands and characterized by heavy erosion, exposed red clay and most prone to fires. Common vegetation found in the grassland sampling areas consisted of the non-native herbaceous plants *Miscanthus floridulus, Pennisetum setaceum, Lygodium microphyllum, Hyptis capitata, Gleichenia linearis, Crotalaria retusa*, and *Arundina graminifolia*. Native plants in the grassland area consisted of *Dicranopteris linearis, Melochia villosissima,* and *Scaevola taccada*. In each sampling site, a 30 m transect was selected that best characterized the type of habitat, and all sampling was performed along the transect and would collect about 3 meters maximum from the transect line if needed. We selected the most abundant plant samples for vegetation beating sheets, and eDNA collections (leaf, flower, and trunk) to understand and compare the overall community. Spider webs were collected randomly along the transit.

### Sample Collections

Two approaches of sampling were compared; (i) bulk DNA metabarcoding that involves direct DNA extraction from samples generated by standard entomological collection methods (malaise trap. beating sheet, white, and yellow pan) and (ii) a non-destructive eDNA approach, where insects were identified via trace amounts of DNA collected from plant surface (flower, leaf, trunk) and spider webs.

**(i) Bulk DNA metabarcoding sampling approaches:** Two malaise traps were deployed at each location, spaced 30 m apart for three days, for a total of 18 malaise traps across Guam. All specimens from malaise traps were collected in 500 mL trap collection bottles and then transferred to 50 mL Falcon tubes. Three each of yellow and white pan traps containing soapy water were placed alternatively every 5 meters along the transect. The insects were collected, and soapy water was replaced daily for three consecutive days, and specimens from the same trap were pooled. Thus, a total of three replicates per pan color resulted from each sampling site, making 27 replicates across Guam. The collected insect samples were strained through a bleach-sterilized metal sieve, rinsed into a 300 ml Nalgene jar containing 95% ethanol, then transferred into 15ml falcon tubes.

Vegetation beating was performed by placing 1 × 1-m white beating sheets under the vegetation and gently agitating the foliage using a 1-m-long PVC pole for timed intervals up to a total beating time of 180 seconds (not including additional time for aspirating and other processing). Arthropods dislodged by the agitation drop onto the beating sheet and are then aspirated into 50 ml Falcon tubes containing 95% ethanol. All collected samples were stored temporarily in −20°C in Guam, then transferred to Evolab at the University of California, Berkeley for analysis.

### Traditional specimen sorting and DNA extraction

Bulk samples from malaise traps, white pans, yellow pans, and beating sheets were screened before DNA extraction. Briefly, the specimens from each sample were spread out in a Petri dish, and then all debris was removed. After cleaning, the specimens were drained, and a wet weight was recorded and then transferred into 15 mL Falcon tubes. However, for bigger individuals above 7 mm, only the legs were included, and the specimen was preserved separately. We used the Qiagen DNeasy Puregene kit (Qiagen, Germany) and followed the modified version for automation described in (Lim et al., 2022), the initial lysis steps were done by simply incubating in 56°C non-destructively, leaving the samples intact for future use.

**(ii) Environmental DNA sampling approaches:** To compare with plant-associated arthropods, the most abundant plants were selected from each site to collect the eDNA samples from the plant surface (flower, leaf, trunk). Intact flowers were opportunistically hand-picked from 5 cm to 200 cm above the forest floor in triplicate per sampling site. Large flowers (> 1 cm petal diameter) were collected in a 50 mL Falcon tube, while small flowers (<1 cm diameter) in a 15 mL tube, both containing 95% ethanol. At each sampling site, 5 to 10 spider webs were collected along the transect or up to two meters from either side of the transect using sterile swab sticks in a 2.0 mL microcentrifuge tube containing Monarch DNA/RNA Protection Reagent (New England Biolabs, Ipswich, MA; catalog #T2011), to capture flying arthropods.

Approximately 200-300g of leaf samples were collected into gallon-sized Ziploc bags along the same 30 m transect from a mixture of abundant plants. The trunk rolling method was adapted from (Valentin et al., 2020). Briefly, a 101mm polyester foam paint roller (Model # HD MR 200-2 4, Linzer Products Corp.) was dampened in 50 mL sterile water and rolled on the trunk and branches. Four strokes of rolling (vertically) were maintained, and each stroke covered ∼1-1.5 m in length. After sampling, paint rollers were stored in Ziploc bags. For the leaf wash and trunk rolling method, 6 replicates were collected, except for N1, M1, and M2 (only three replicates). Thus, in total, 45 samples were collected for each leaf wash and trunk rolling across Guam.

### eDNA Sample Processing

For leaf wash and trunk rolling samples, washing with double-distilled water and then subsequent filtration were employed. 750 mL of double-distilled water was added to each Ziploc bag containing leaves and trunk rollers. The samples were subjected to brief shaking for 10 seconds, followed by a 15-minute settling period to facilitate the transfer of eDNA into the water. Subsequently, the water was pre-filtered using sterile coffee filters to eliminate large debris, minimizing the risk of clogging during downstream filtration processes. 500 ml of water was directly filtered through a 250ml Nalgene™ Sterile Analytical Filter equipped with 0.45 μm cellulose nitrate membranes (Thermo Fisher Scientific, USA) by using a Cole-Palmer Masterflex® peristaltic pump (Cole-Parmer, USA), powered by an electric drill (Laramie et al., 2015). Following filtration, the filter papers were carefully removed using sterilized forceps and transferred into 2.0 mL microcentrifuge tubes containing 95% ethanol. All samples were stored temporarily in −20°C at the Center for Environmental Management of Military Lands lab, Guam, and then transferred to Evolab at the University of California, Berkeley, for further analysis.

In total 313 samples were collected, where 117 samples were from four bulk DNA metabarcoding methods [malaise trap: n = 18, beating sheet: n = 45, white pan: n = 27, yellow pan: n = 27]; and 196 samples from four eDNA metabarcoding methods [leaf wash: n = 45, trunk rolling: n = 45, spider web: n = 79, and flower: n = 27].

All necessary measures were taken to control field contamination, briefly (i) sterile nitrile gloves were used during collection and processing, (ii) all collection bags, falcon tubes, collection vials were new and sterile, (iii) complete separate set of field kits (gloves, tissues, markers, Ziploc bags) for eDNA and bulk DNA metabarcoding were used, (iv) after collection all eDNA samples were stored in a Styrofoam box with ice to prevent degradation.

### Environmental DNA extraction

The eDNA extractions were performed in a separate workspace dedicated to low DNA concentration at the Evolab, University of California, Berkeley. The eDNA from flowers, spider webs, trunk rolling and leaf washes was extracted using the DNeasy Blood and Tissue Kit (Qiagen, Germany), with the following modifications for each sample: (i) in the case of flowers, a centrifugation at 8000 rpm for 10 min was done to form the pellet of suspended residues in ethanol, then the ethanol was discarded and samples were air dried in a hot air-oven (60°C) for 4h before adding the lysis buffer. In each sample, 540 μL ATL (tissue lysis) buffer and 60 μL proteinase K were added to incubate the samples at 56°C for an overnight lysis. (ii) Leaf wash and trunk rolling filter papers were cut into two halves, where one half of the filter paper was stored in the same collection tube for future use, and the other half was air dried in a hot air oven for 4 h to remove ethanol for further DNA extraction process. After that, 180 μL ATL buffer and 20 μL proteinase K were added to incubate the samples at 56°C for an overnight lysis. (iii) Spider web samples were directly incubated (as they were preserved in Monarch^®^ DNA buffer) at 56°C for an overnight lysis with 180 μL ATL buffer and 20 μL proteinase K.

After the lysis, the standard protocol of DNeasy Blood and Tissue Kit (Qiagen, Germany) with some modification (as described in Banerjee et al., 2025) was followed except 600 μL AL buffer for flower samples. The quality and quantity were checked using nanodrop spectrophotometer (Implen, Germany) and a qubit fluorometer (Thermo Fisher Scientific, USA). All DNA extracts were stored at −20°C for further analysis.

### PCR amplification, library preparation, and sequencing

All DNA samples (except spider webs) were amplified using two universal degenerate primer pairs amplifying the mitochondrial COI (mt-DNA) barcoding region; (i) mlCOIintF (5′--GGWACWGGWTGAACWGTWTAYCCYCC--3′; Leray et al., 2013) and Fol--degen--rev (5′--TANACYTCNGGRTGNCCRAARAAYCA--3′; Yu et al., 2012); for 313-bp amplicon (hereafter referred to as MCO) and (ii) LCO1490 (5′ GGTCAACAAATCATAAAGATATTGG 3′; Folmer et al., 1994) and CO1 CFMRa (5′ GGWACTAATCAATTTCCAAATCC 3′; Jusino et al., 2019); for 180-bp amplicon (hereafter refer as ANML). For spider webs, we used only the ANML primer. A two-step PCR protocol was employed for library preparation. The initial PCR was performed in triplicate with a 6-bp inline barcode combination. Each 10 μL reaction mixture comprises 5 μL of 2× QIAGEN Multiplex PCR Master Mix (Qiagen, Germany), 1 μL of DNA template, 0.5 μL of each 10 μM forward and reverse primer, 1 μL of Q solution, and 2 μL of nuclease-free water. PCR parameters were 95°C for 15 mins, 35 cycles of 94°C for 30s, 55°C for 90s, 72°C for 90s and a final extension of 72°C for 10 min. Pooling of all subsamples was done based on band intensity observed during 2% agarose gel electrophoresis with Gel Red (Fisher Scientific, USA). The second PCR (indexing PCR) was used to attach sequencing adapters and 8-bp dual indexes to each sample, ensuring a minimum 2 bp difference between indexes (Lange et al., 2014). The PCR parameters were the same as before, except only five cycles instead of 35 cycles. All field, filtration, extraction, and PCR blanks were used as negative controls. After a final screening of agarose gel electrophoresis, all PCR products were cleaned using 1.5x SPRI beads (AMPure XP, USA), and quality control was done using Invitrogen qubit 3.0 fluorometer (Thermo Fisher Scientific, USA) and Agilent Bioanalyzer 2100 system (Agilent Technologies, USA) and then pooled in equal amounts into a single tube. Finally, all samples were sequenced using NextSeq 2000 P2 600 Cycle (300PE) with 30% PhiX at QB3 Genomics Facility, University of California, Berkeley.

### Bioinformatic analysis

After sequencing, all reads were demultiplexed by the QB3 Genomics Facility, University of California, Berkeley, and raw sequences were provided. PEAR (Zhang et al., 2014). was used to merge the demultiplexed reads with a minimum overlap of 50 bp and a quality threshold of 20. The FastX toolkit (Gordon, & Hannon, 2010) was used to convert reads to FASTA files after quality filtering for at least 90% of bases exceeding Q30. Sequences of primers from both ends were removed in UNIX. USEARCH was used to dereplicate the reads, and finally, zero-radius OTUs (zOTUs) were prepared using the unoise3 command (Edgar, 2010, 2016). After error removal, BLASTN was used to blast all zOTUs against the GenBank nucleotide database (downloaded January 2024).

After this, a stringent taxonomic filtration was done on the following criteria to eliminate all potential contamination: (i) (phylum= “Arthropoda”, percentage of identification>= 80%, sequence length > 170 for ANML primer, and sequence length > =280 for MCO); (ii) samples with low sequence depth of reads (<5000 reads) were discarded; (iii) OTUs with total reads < 50 were discarded; (iv) OTUs detected in negative control with > 50 reads in blank were removed from samples. Taxonomic assignments were done based on a similar threshold of 90% for family, 95% for genus, and over 98% for species. A huge portion of OTUs could not be identified to genus or species level, so we restricted our analyses to the observed OTU level.

All data analyses and statistics were performed using R version 4.3.3 (R Core Team, 2020). Samples were rarefied using the rarefy() function in vegan, based on the minimum read count across all samples with 999 permutations (Schloss, 2024). To visualize differences in taxonomic composition between method groups (bulk DNA and eDNA metabarcoding), within methods (malaise trap, beating sheets, white pan, leaf wash, flower, etc.), and among habitats (intact limestone forest, degraded forest, and grassland), alpha diversity was visualized based on observed OTU and species richness using ggplot2 (Hamilton & Ferry, 2018); and a non-metric multidimensional scaling (NMDS) analysis was performed using Jaccard similarity matrices with the metaMDS function (k = 2, trymax = 999) in the vegan R package (Oksanen et al., 2019). OTU and species composition overlap between method groups was visualized by eulerr (Larsson & Gustafsson, 2018). The difference in detection effectiveness and overlap between methods (malaise trap, beating sheet, white pan, leaf wash, flower, etc.), was visualized using the Upset plot. The overall differences between methods were tested using PERMANOVA (adonis2() in vegan, Jaccard dissimilarity, 999 permutations); pairwise differences between the method group and habitat were tested using the pairwise Wilcoxon’s test.

## Results

### Overall amplifications

Both primer pairs (ANML and MCO) were effective in recovering taxonomic diversity across all collecting approaches and habitats. In the case of the ANML primer, 51,633,216 reads were generated from 308 samples, making 167,640 (±7524) mean reads per sample. After quality control and taxonomic assignments, 25,698,361 DNA reads from 239 samples were found to be associated with 6188 OTUs from diverse groups (Figure 2). Among all OTUs, the majority were assigned to Arthropoda, followed by Rotifera, Mollusca, Discosea, and Tubulinea (Figure 2). After filtering Arthropoda, we retained 5,176 OTUs across 239 samples collected using the different methods (Figure S1). In total, five samples failed to amplify at the PCR stage, and 69 samples failed to pass the post screening test.

In the case of the MCO primer, a total of 36,228,227 DNA reads were generated from 215 samples, making 168,503 (±8392) mean reads per sample. After quality control and taxonomic assignments, 19,635,935 DNA reads from 188 samples were found to be associated with 15,476 OTUs from diverse groups (Figure 2). The majority of the OTUs were detected from Arthropoda, followed by Ascomycota and Basidiomycota (Figure 2). After filtering Arthropoda, we retained 8825 OTUs across 188 samples collected using different methods (Figure S2). Here, 19 samples at the PCR stage and 27 samples failed to pass the screening test (described in the method section). Although MCO detected higher OTUs than ANML, both primers showed similarities in the proportion of arthropod taxa detected (Figure 2). Predictably, only 68 OTUs from ANML and 55 OTUs from MCO could be identified to genus or species level, likely due to limitations in the reference database.

Both primer sets yielded broadly comparable results (Figure 2). Because of differences in amplification efficiency and sample coverage, we analysed the OTU results separately rather than merging them. As the ANML primer amplified the majority of samples, it was used for downstream analyses; corresponding results from the MCO primer are provided in the supplementary materials.

### Bulk DNA metabarcoding vs eDNA metabarcoding

All the collecting approaches for bulk DNA metabarcoding detected greater OTU diversity compared to the eDNA metabarcoding (**Figure 3A**). The detection rate significantly differs between bulk DNA and eDNA metabarcoding based on the overall observed OTU (p < 0.001), where bulk DNA metabarcoding detects a larger number of taxa compared to eDNA metabarcoding (**Figure 3B**). Among the bulk DNA metabarcoding methods, the beating sheet detected the highest OTUs, followed by the white pan, yellow pan, and malaise trap. In the case of eDNA metabarcoding, trunk rolling detected the highest number of OTUs, followed by leaf wash, flowers, and spider webs (**Figure 3A**). The Venn diagram shows bulk DNA metabarcoding significantly dominating and contributing a large portion of unique and overlapping detection (**Figure 4A**). Similar results were also noted in the Upset plot (**Figure 4B**). Interestingly, 47% of OTUs were detected across both bulk DNA and eDNA approaches, where bulk DNA metabarcoding uniquely detected 40%, and eDNA detected 13% (**Figure 4A**). Although eDNA methods detected fewer taxa compared to bulk DNA metabarcoding, eDNA covers almost 60 percent of the overall detection. The data show that a combination of all methods maximizes arthropod detection (**Figure 4B**).

### Ecological insights: Multivariate patterns across methods and habitats

Alpha diversity based on observed OTUs showed significant differences among the three habitats (**Figure 5**), although the results varied between primer pairs (Figure S4). The NMDS plot shows that the broad categories of bulk DNA and eDNA metabarcoding methods were clustered separately (**Figure 6**). Three eDNA methods – leaf wash, spider web, trunk rolling – clustered together, whereas the flower community was distinct from others (**Figure 6**). In bulk DNA metabarcoding methods, white pan and yellow pan were highly clustered together. Communities in malaise traps were widespread and showed resemblance with beating sheet, white pan, yellow pan, and flower.

The ANOSIM results indicated that the arthropod communities detected by different sampling methods are significantly different in composition (R = 0.632, p = 0.001), with much greater dissimilarity between methods than within any single method. Arthropod communities showed only weak but statistically significant differences among habitats (R = 0.142, p = 0.001). The PERMANOVA results revealed that the sampling method was the primary factor driving differences in arthropod community composition, explaining approximately 17% of the total variation (R² = 0.169, F = 4.85, p = 0.001). Habitat type also had a statistically significant but much smaller effect, accounting for only about 3.8% of the variation (R² = 0.038, F = 2.89, p = 0.001). When both factors were considered together, they explained about 20.4% of the variation (R² = 0.204, F = 4.52, p = 0.001). Pairwise PERMANOVA comparisons demonstrated that most sampling method pairs differed significantly in arthropod community composition, with p-values below 0.001 in nearly all cases (**Table 2**). The biggest differences were observed between the beating sheet and several other methods (e.g., white pan, yellow pan), each explaining 9.6–13.5% of variation (R²). Some comparisons, such as white pan vs yellow pan, showed weaker or borderline significant differences (p = 0.053), suggesting more similarity between these methods (**Table 2**).

**Table 2.** Pairwise PERMANOVA results comparing arthropod community composition across different sampling methods. The table shows degrees of freedom (Df), sum of squares (SumOfSqs), effect size (R²), F-statistic (F), p-values, and significance levels for each pairwise comparison. Most method pairs differed significantly (p < 0.001), indicating that sampling method strongly influences community composition detected in this study.

| Comparison | Df | SumOfSqs | $R^2$ | F | p-value | Significance |
| --- | --- | --- | --- | --- | --- | --- |
| Beat Sheet vs White Pan | 1 | 3.0592 | 0.13339 | 8.158 | 0.001 | *** |
| Beat Sheet vs Yellow Pan | 1 | 3.2943 | 0.13458 | 8.7082 | 0.001 | *** |
| Beat Sheet vs Malaise Trap | 1 | 1.7804 | 0.09626 | 4.8996 | 0.001 | *** |
| Beat Sheet vs Leaf Wash | 1 | 1.9456 | 0.09935 | 5.7361 | 0.001 | *** |
| Beat Sheet vs Trunk Rolling | 1 | 2.5406 | 0.09638 | 6.3994 | 0.001 | *** |
| Beat Sheet vs Flower | 1 | 1.3116 | 0.09261 | 3.6741 | 0.001 | *** |
| White Pan vs Yellow Pan | 1 | 0.5874 | 0.03003 | 1.3931 | 0.053 | . |
| White Pan vs Malaise Trap | 1 | 1.5637 | 0.09732 | 3.7735 | 0.001 | *** |
| White Pan vs Leaf Wash | 1 | 2.8913 | 0.15785 | 7.6846 | 0.001 | *** |
| White Pan vs Trunk Rolling | 1 | 1.5557 | 0.06716 | 3.5277 | 0.001 | *** |
| White Pan vs Flower | 1 | 0.9546 | 0.08233 | 2.2429 | 0.001 | *** |
| Yellow Pan vs Malaise Trap | 1 | 1.7310 | 0.09866 | 4.1596 | 0.001 | *** |
| Yellow Pan vs Leaf Wash | 1 | 3.0642 | 0.15476 | 8.056 | 0.001 | *** |
| Yellow Pan vs Trunk Rolling | 1 | 1.6876 | 0.06858 | 3.829 | 0.001 | *** |
| Yellow Pan vs Flower | 1 | 0.9625 | 0.07454 | 2.2551 | 0.001 | *** |
| Malaise Trap vs Leaf Wash | 1 | 1.7539 | 0.12510 | 4.8615 | 0.001 | *** |
| Malaise Trap vs Trunk Roll | 1 | 1.4326 | 0.07206 | 3.2613 | 0.001 | *** |
| Malaise Trap vs Flower | 1 | 0.9662 | 0.11438 | 2.3247 | 0.001 | *** |
| Leaf Wash vs Trunk Rolling | 1 | 2.4989 | 0.11426 | 6.192 | 0.001 | *** |
| Leaf Wash vs Flower | 1 | 1.3033 | 0.13428 | 3.7225 | 0.001 | *** |
| Trunk Rolling vs Flower | 1 | 0.8383 | 0.05435 | 1.8392 | 0.003 | ** |

## Discussion

### Overall amplification: Differences between primer pairs

Differences in taxonomic recovery between the ANML and MCO primers are due to the degeneracy of both primers. The higher OTU richness observed with MCO may be partly attributable to its highly degenerate nature. Low species-level identifications in both primer sets underscore the limitations of current reference databases, particularly for island ecosystems like Guam, where many taxa remain undescribed or unrepresented in barcode libraries. These results highlight the need for expanded reference datasets to improve molecular resolution in undersampled systems. Besides, most of the failed MCO amplifications were from flower and beating sheet samples, whereas in the case of ANML primer spider web was the main contributor of failed samples. This pattern likely reflects a technical issue rather than primer bias. Additionally, failure of 70% spiderweb eDNA suggests more methodological work on developing new sources.

### Choice of method: eDNA and/or bulk DNA metabarcoding

Non-destructive eDNA metabarcoding contributed significantly to the detection of taxa, reinforcing its value as a method for monitoring biodiversity in sensitive environments. Bulk DNA metabarcoding consistently recovered a higher number of OTUs compared to eDNA metabarcoding (**Figures 3 and 4**). These results align with the previous findings that bulk-sample DNA generally contains higher concentrations of intact target DNA, resulting in greater sequencing depth and broader taxonomic coverage (Hajibabaei et al., 2019; Roger et al., 2022). Despite the overall higher performance of bulk DNA metabarcoding, eDNA methods contributed 60% of overall detections (Figure 4). This validates the importance of eDNA as an alternative method that can capture transient taxa (e.g., from aerial deposition or rare encounters) and remnant trace biological material (e.g., shed exoskeletons, frass, casings) that may be absent from physical collections (Weber et al., 2024a). Thus, eDNA and bulk DNA metabarcoding can complement each other, underscoring the value of both approaches to improve biodiversity inventories and reduce false absences (Weber et al., 2024b). Overall, eDNA and bulk DNA metabarcoding both detected large portions of the arthropod diversity, with an overlap of 47% across all collecting approaches. Moreover, the sensitivity of eDNA allows trace amounts to be recovered from the environment, potentially enabling capture of additional diversity relative to established trapping methods.

### Ecological insights: Multivariate patterns across methods and habitats

Our multivariate analyses showed that the sampling approach was the dominant driver in the detection of differences in arthropod diversity, explaining ∼17% of the observed variation. However, habitat type accounted for only ∼4% of the variation. This strong methodological effect is in line with previous multi-method biodiversity surveys (Roger et al., 2022; Kestel et al., 2023; Weber et al., 2024a; Jones et al., 2025), emphasizing that the choice of collection method profoundly shapes observed community structure. The clustering patterns in NMDS ordinations reflected functional similarities between certain methods (e.g., white and yellow pan), while others (e.g., trunk rolling and flower) captured more distinct assemblages (**Figure 6**). Notably, some eDNA methods (e.g., leaf wash, spider web, trunk rolling) clustered closely with bulk-sample methods (beating sheet and malaise traps), suggesting partial overlap in the assemblages they capture, potentially due to shared microhabitats as we tried to compare (Figure 6, Table 1). The high number of unique taxa detected by individual methods in the Upset analysis and the significant pairwise PERMANOVA differences between most methods suggest that no single approach can adequately represent total arthropod diversity (**Figure 4**, **Table 2**). This finding reinforces the need for method integration in large-scale monitoring schemes, particularly in heterogeneous landscapes where taxa are differentially detectable across substrates. However, a single sampling strategy can be enough to understand the species interactions (flower for monitoring pollinators) or targeting particular species for monitoring.

The finding of significant differences in observed OTUs among the three habitats (**Figure 5**) is further supported by the NMDS analysis (**Figure 6**), suggesting that while both sampling method and habitat influence arthropod community structure, including multiple sampling approaches is important to maximize assessment of diversity.

However, habitats may also have significant effects, which may be overlooked because of reference databases. Thus, a stronger reference database could help in identifying OTUs to genus or species level, which potentially helps in understanding the effect of habitats on species distribution. Thus, building a reference database for more species-specific monitoring may be the ideal next step for Guam.

## Conclusions

Our results indicate that both DNA and eDNA metabarcoding methods are fast and effective for recovering a broad spectrum of the terrestrial arthropod diversity in a remote island ecosystem. eDNA recovered a large proportion of the taxa recovered by bulk DNA metabarcoding. While bulk DNA metabarcoding approaches recovered higher OTUs per sample than eDNA, the sampling method was the dominant source of variation in explaining arthropod diversity. The detection efficiency across habitat types suggests that both bulk DNA and eDNA metabarcoding are suitable for rapid biodiversity monitoring when multiple collection methods are used, even when complete DNA reference libraries are limited. Further, eDNA metabarcoding can be a non-destructive alternative in sensitive habitats where insect capture is undesirable and can provide additional data for particular species interactions.

## Supporting information

Arthropoda, we retained 5,176 OTUs across 239 samples collected using the different methods (Figure S1).

## Author Contributions

Kenneth Puliafico, Haldre Rogers, Samantha Al-Bayer, Jerilyn Calaor, Natalie Graham, George Roderick, and Rosemary Gillespie designed the project. Samantha Al-Bayer, Jerilyn Calaor, Natalie Graham, and Kenneth Puliafico conducted the fieldwork. Pritam Banerjee, Samantha Al-Bayer, and Sven Weber performed laboratory work for eDNA metabarcoding. Pritam Banerjee and Sven Weber carried out bioinformatic processing of the samples. Pritam Banerjee conducted the data analysis and prepared the first draft of the manuscript with input from all authors. All authors contributed to and approved the final manuscript.

## Acknowledgements

The authors thank Paul Krushelnycky and Dimitrios Petsopoulos for their initial contributions to this project. They also thank the sequencing facilities at UC Berkeley for their continuous support: the QB3 Genomics Facility (see https://qb3.berkeley.edu/facility/genomics/cite/) and the UC Berkeley DNA Sequencing Facility (see https://ucberkeleydnasequencing.com).

## Ethics Statement

The authors have nothing to report.

## Consent

The authors have nothing to report.

## Data Availability Statement

The data supporting this study are available for peer review at Dryad. The dataset will be made publicly available upon acceptance of the manuscript.

---

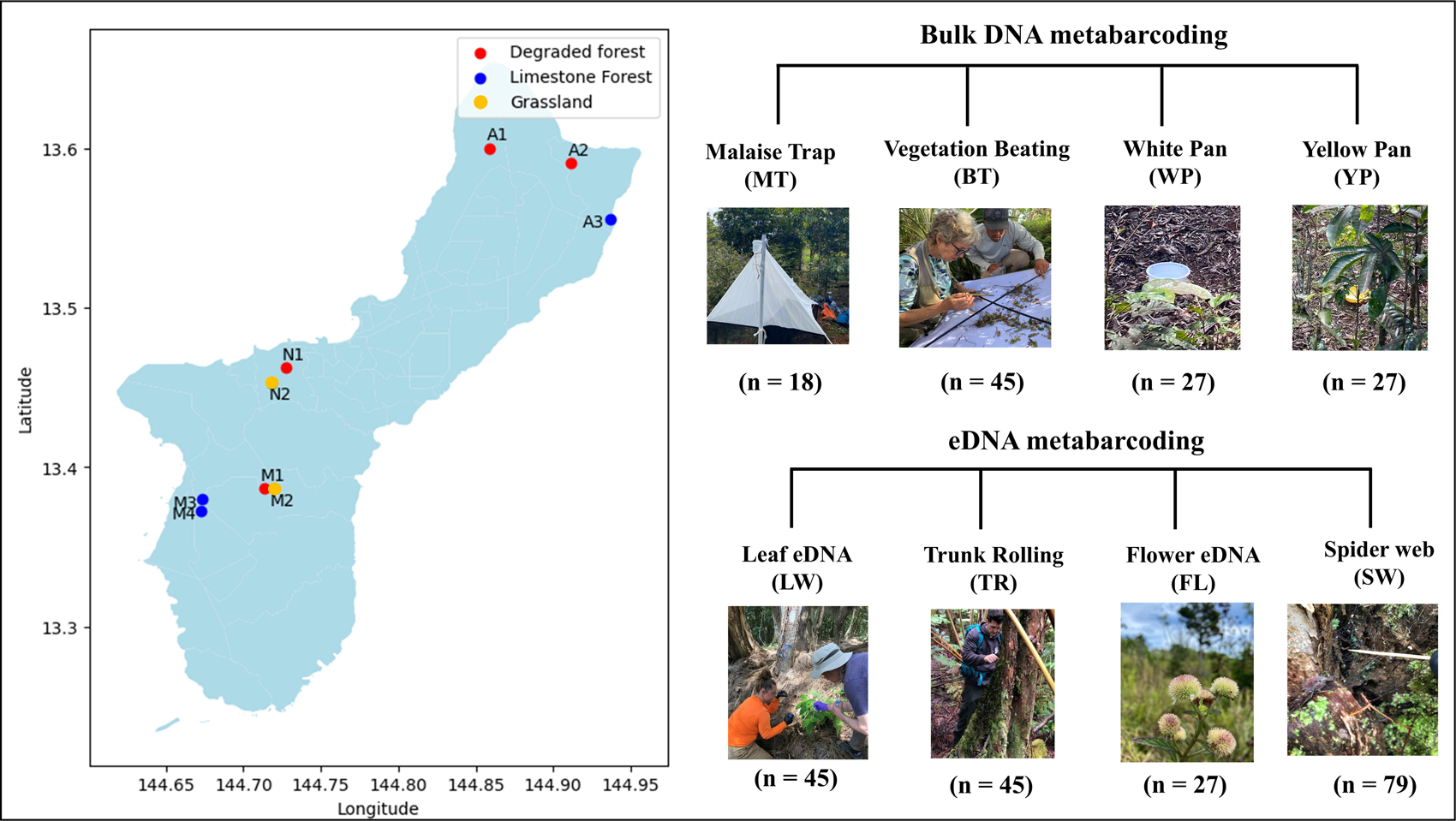

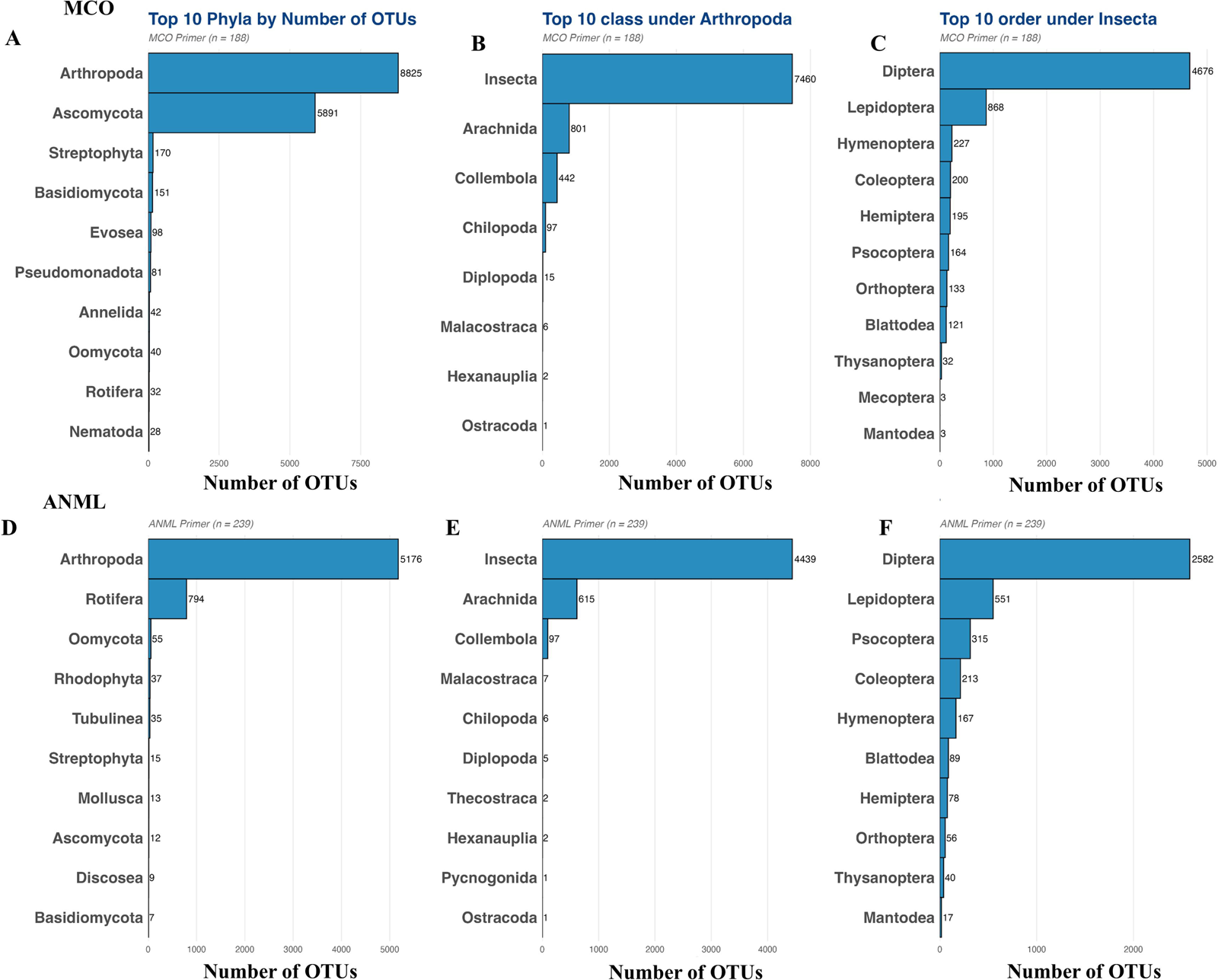

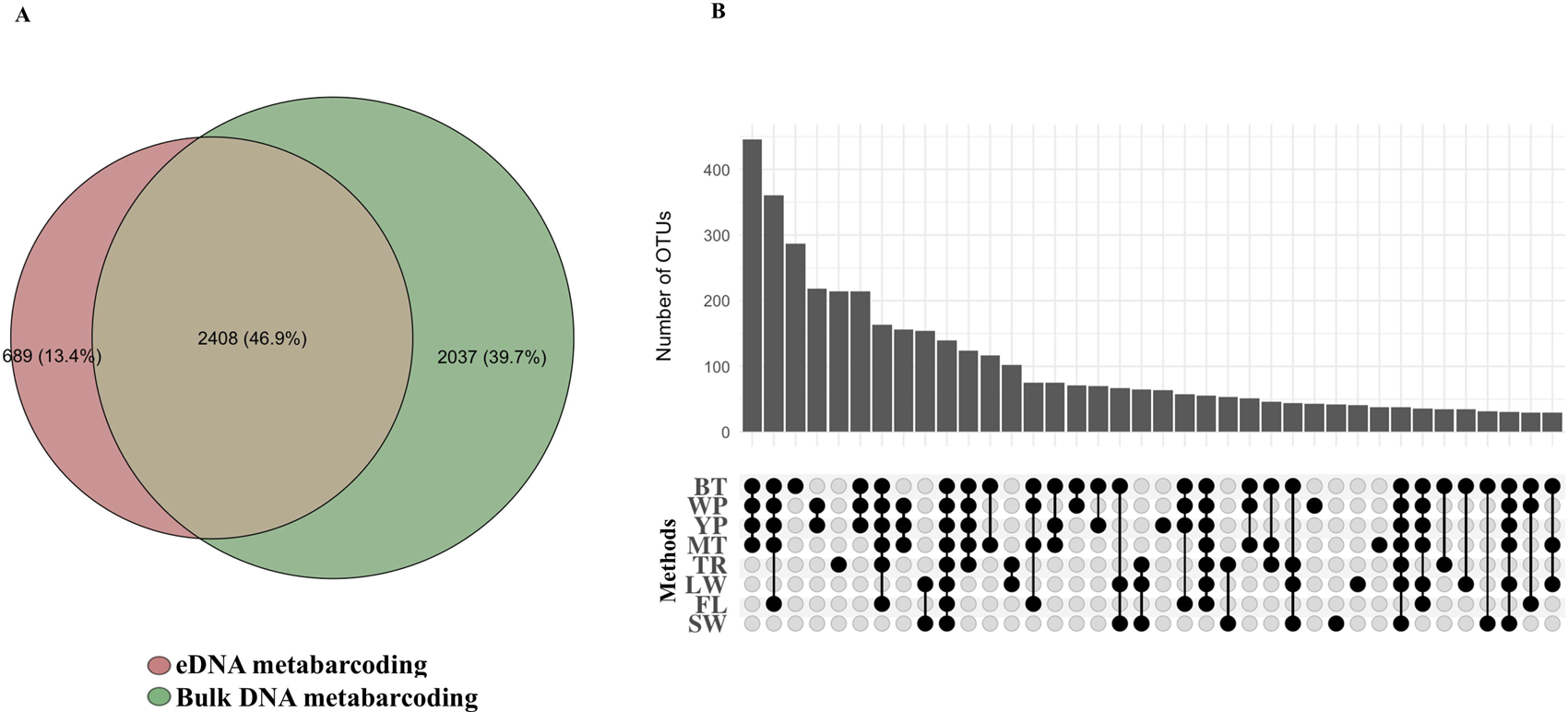

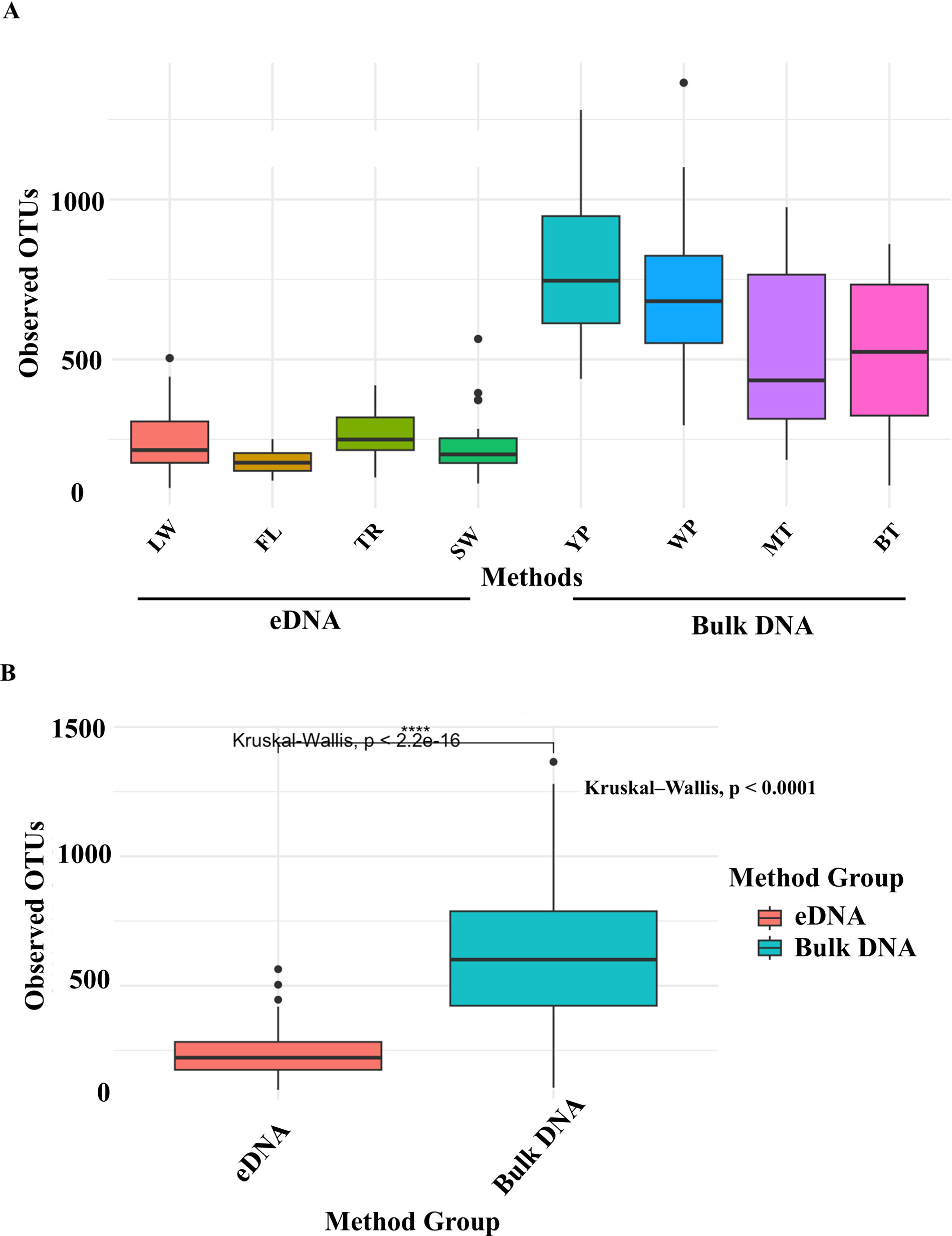

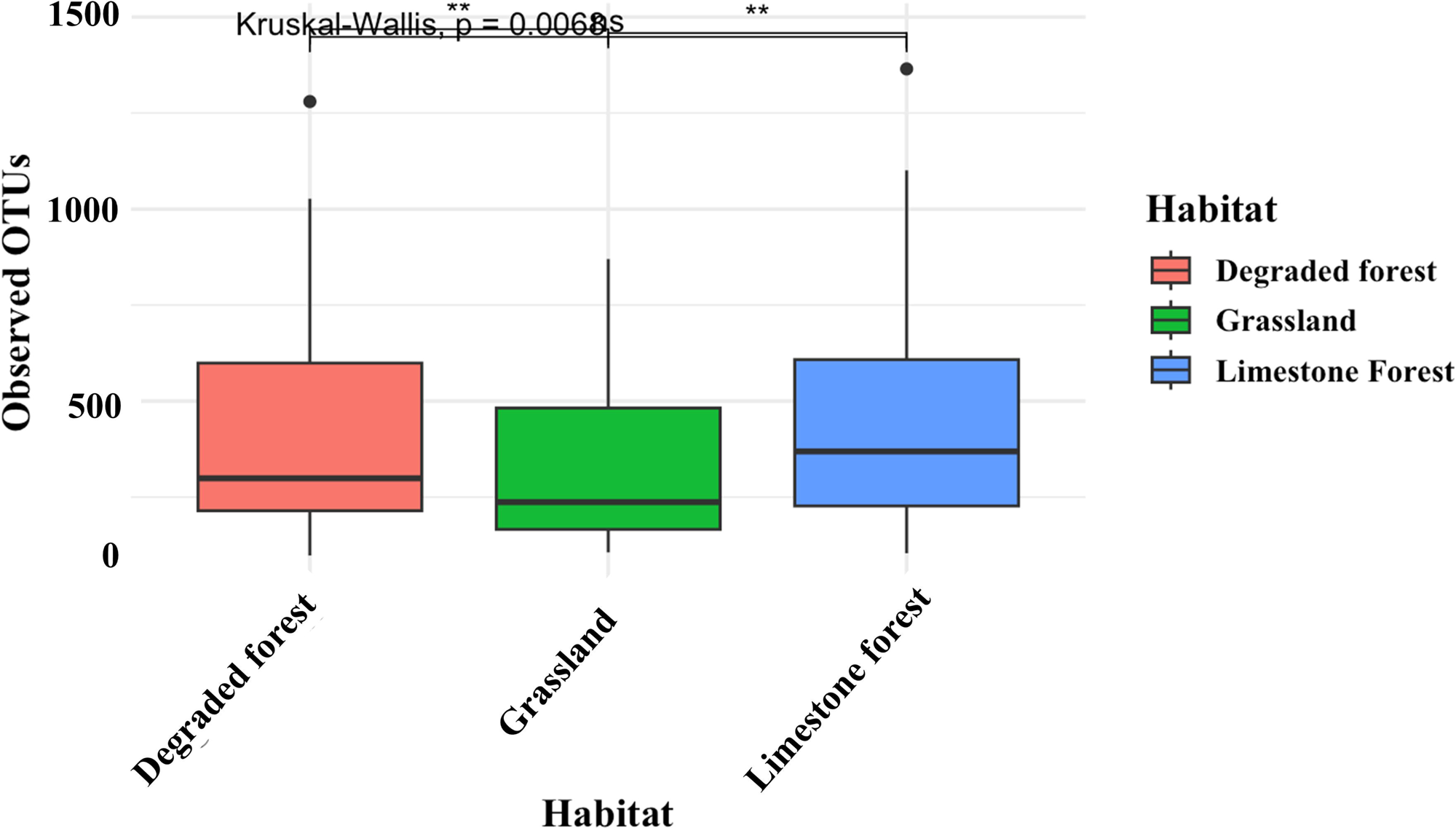

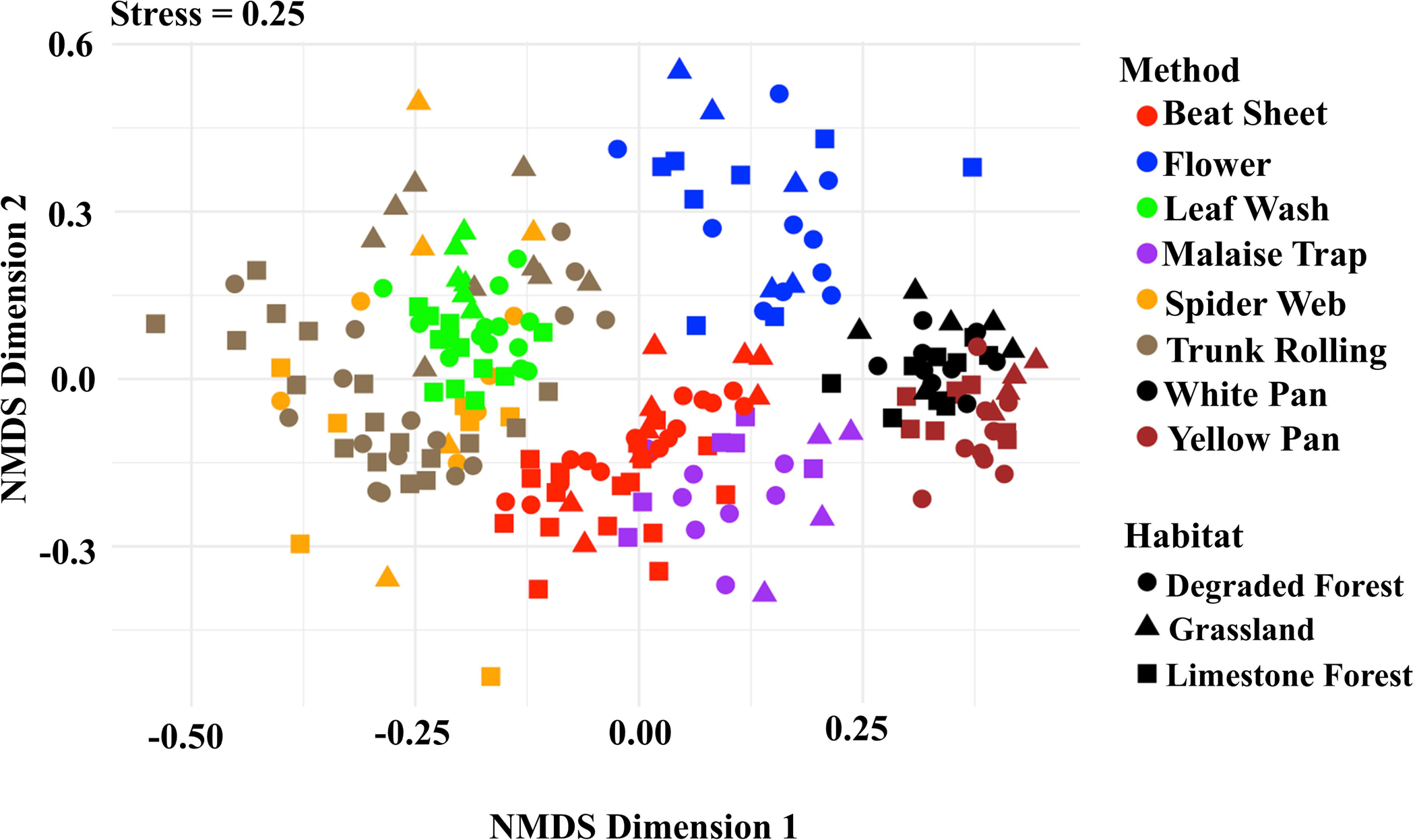

