## Supplementary material for "Comparison of environmental DNA and bulk DNA metabarcoding for assessing terrestrial arthropod diversity across three habitat types on Guam": Arthropoda, we retained 5,176 OTUs across 239 samples collected using the different methods (Figure S1).

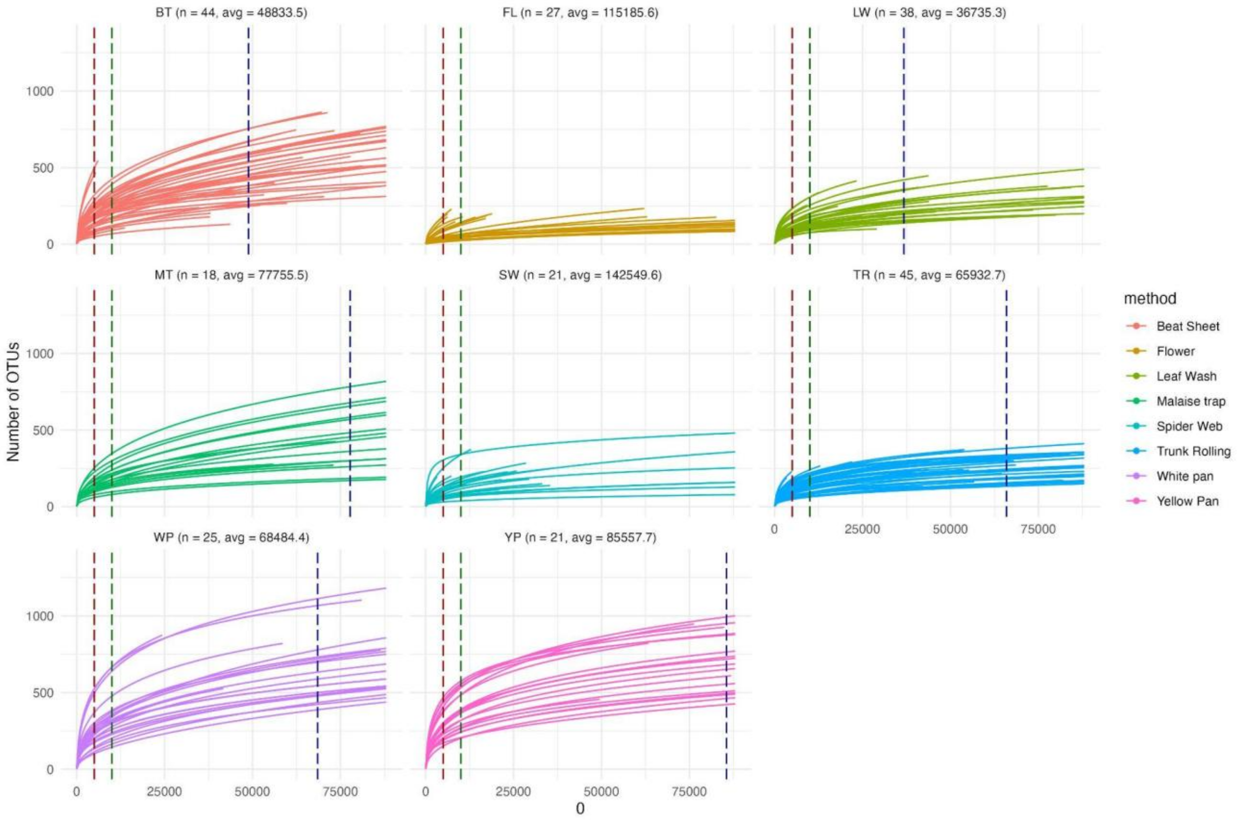


**Figure S1.** Rarefaction curve of the primer ANML after filtering for Arthropoda. We retained 5,176 OTUs across 239 samples collected using different methods, including malaise trap (n = 18, average reads = 77755), beating sheet (n = 44, average reads = 48833), white pan (n = 25, average reads = 68484), yellow pan (n = 21, average reads = 85557), leaf wash (n = 38, average reads = 36735), trunk rolling (n = 45, average reads = 65812), spider web (n = 21, average reads = 142549), and flower (n = 27, average reads = 115185). Five samples failed to amplify at the PCR stage and 69 samples failed to pass the screening test for ANML primer due to total reads > 5000. Among the lost samples, 58 samples were from spider web, 7 samples from leaf wash, 6 from yellow pan, 2 from white pan, and 1 from beating sheet.

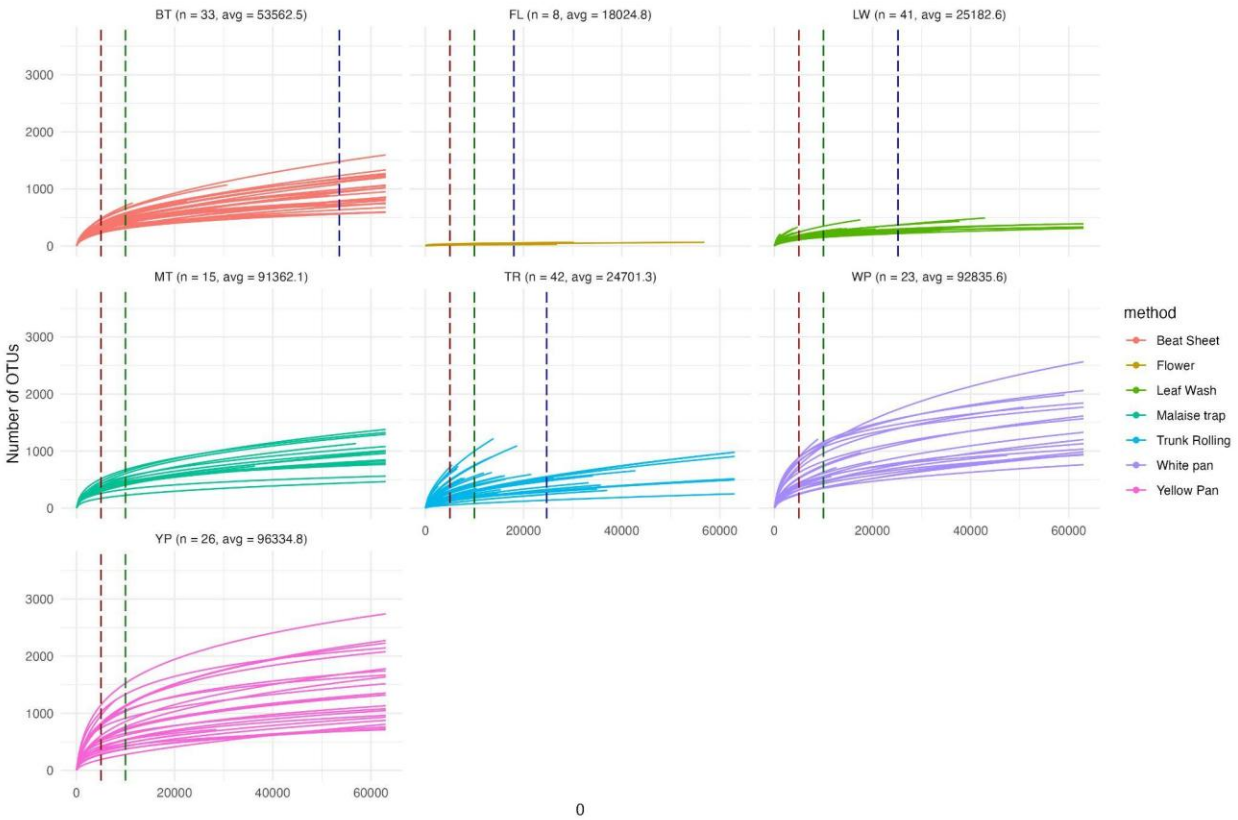


**Figure S2.** Rarefaction curve of the primer MCO after filtering for Arthropoda. We retained 8825 OTUs across 188 samples collected using different methods, including malaise trap (n = 15, average reads = 91,362), beat sheets (n = 33, average reads = 53,562), white pans (n = 23, average reads = 92,835), yellow pans (n = 26, average reads = 96,334), leaf washes (n = 41, average reads = 25,182), trunk rollings (n = 42, average reads = 24,701), and flowers (n = 8, average reads = 18,024) (Figure S2). Here, 19 samples failed to amplify, and 27 samples failed to pass the screening test for the MCO primer due to total reads > 5000. Among the lost samples, 19 samples were from flowers, 12 samples from beating sheet, 4 from white pan and leaf wash: 3 from malaise trap and trunk rolling, and one from yellow pan.

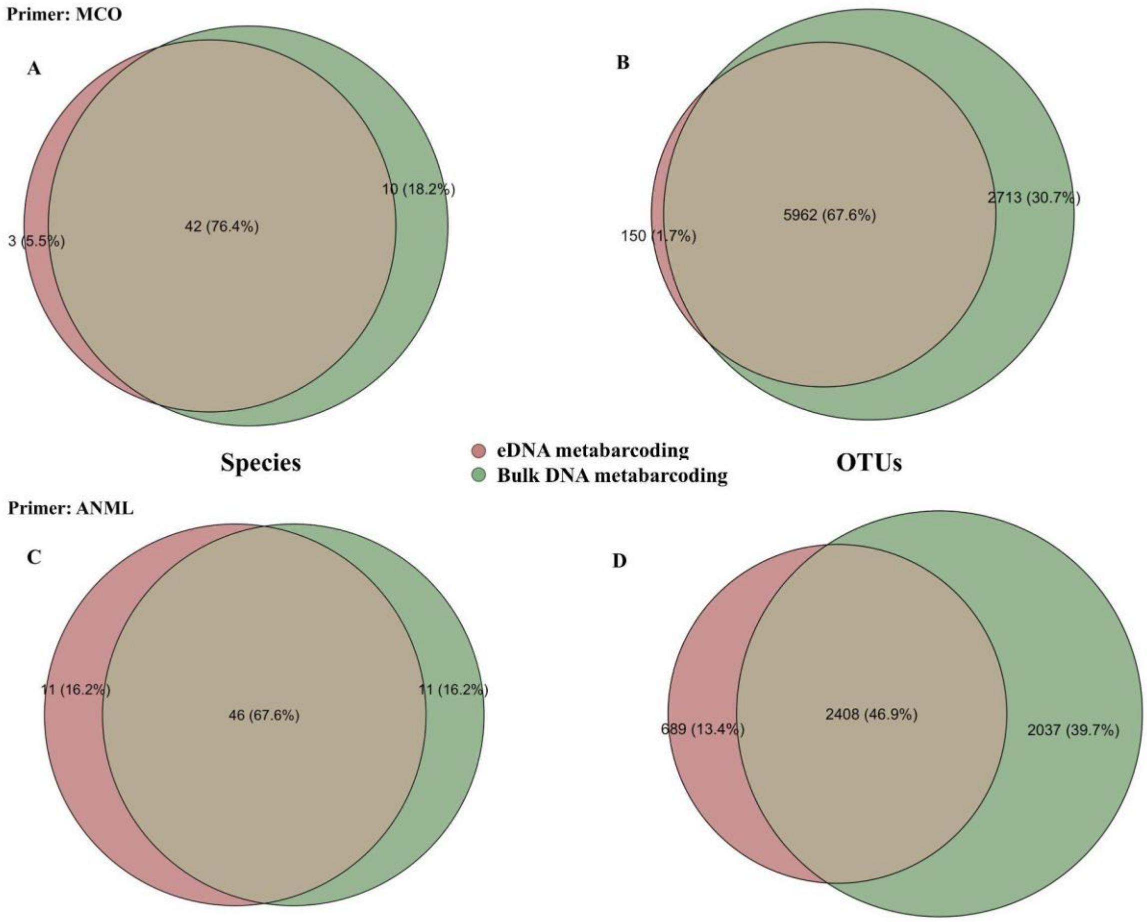


**Figure S3.** Venn diagrams showing the overlap in species and OTU detections obtained with two different primers, MCO (A, B) and ANML (C, D), for both bulk DNA metabarcoding and eDNA metabarcoding.

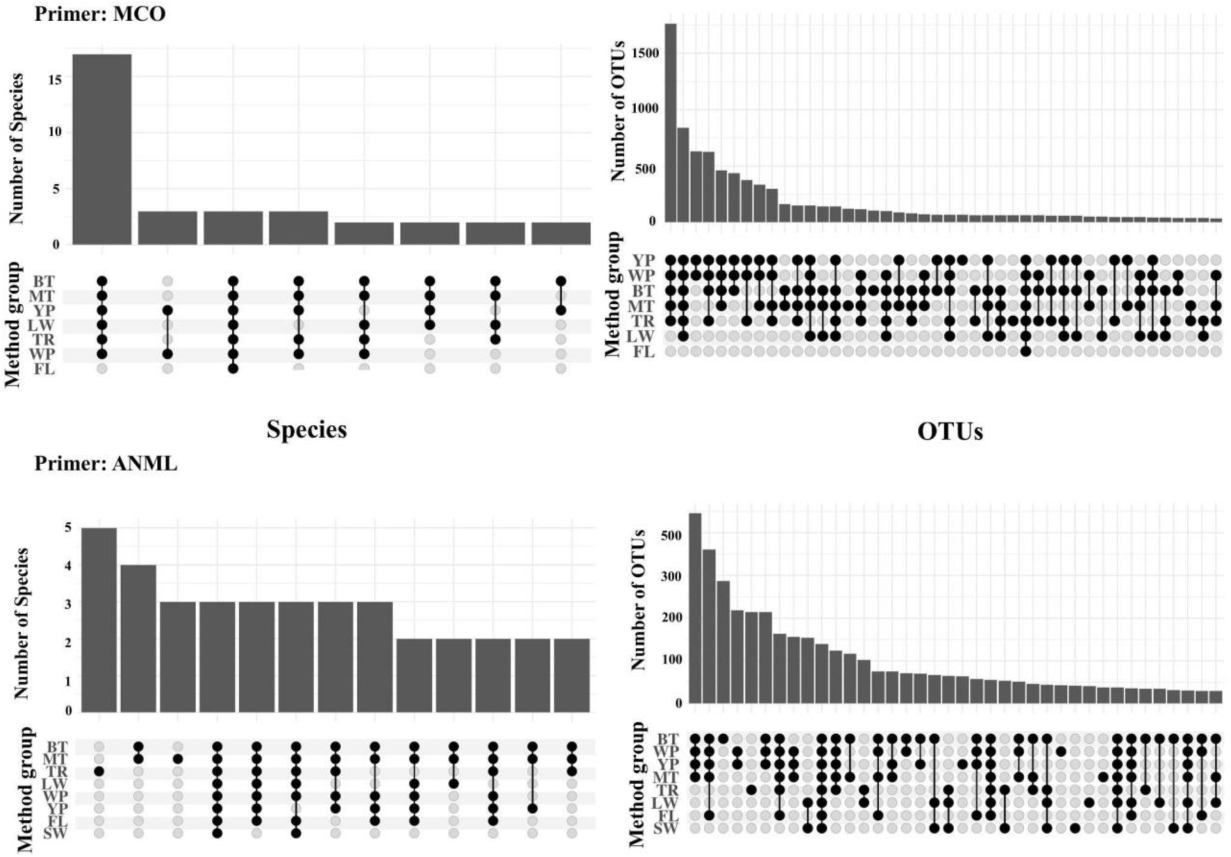


**Figure S4.** Upset plots showing the cumulative detection of species (left panels) and OTUs (right panels) across different sampling methods using two primer sets: MCO (top row) and ANML (bottom row). Bars represent the number of species or OTUs detected, while connected dots indicate the intersections or cumulative detection among methods. Note: MT: malaise traps; BT: beating sheets; WP: white pan; YP: yellow pan; FL: flower; LW: leaf wash; TR: trunk rolling; and SW: spider webs.

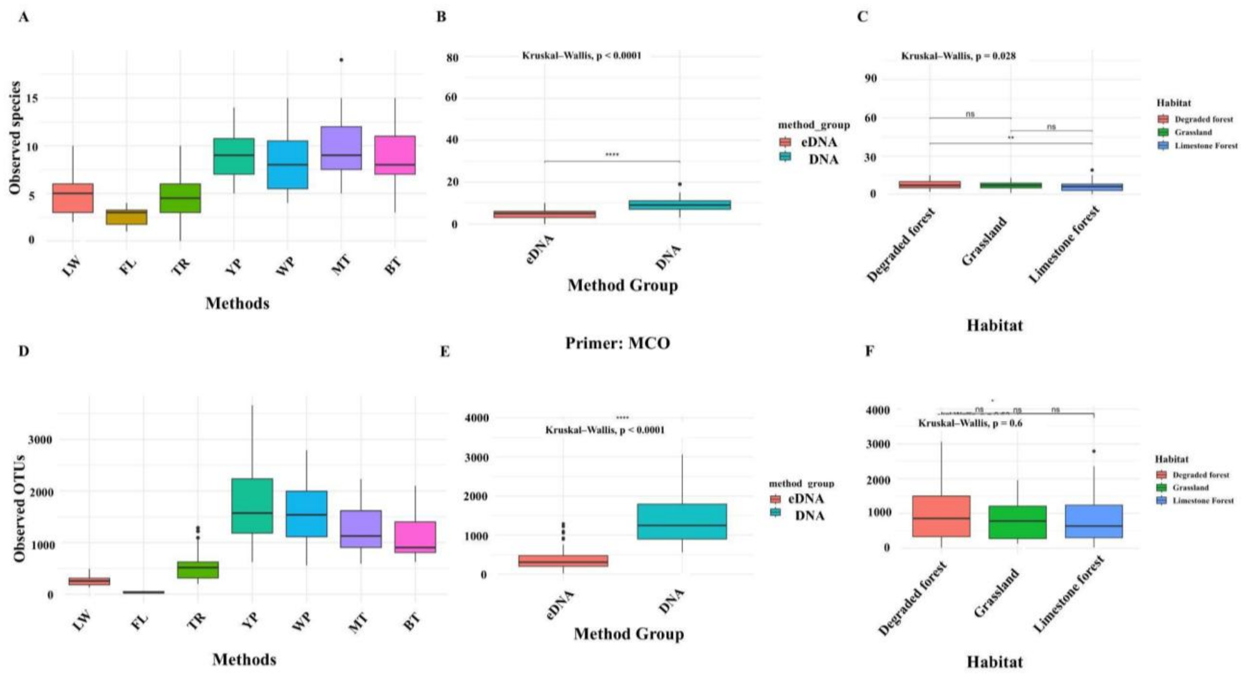


**Figure S5.** Alpha diversity of species (A–C) and OTUs (D–F) detected using the MCO primer across different methods, method groups, and habitats. (A, D) Boxplots showing the number of observed species and OTUs across individual sampling methods (MT: malaise traps, BT: beating sheets, WP: White Pan, YP: Yellow Pan, FL: flower, LW: leaf wash, and TR: trunk rolling,). (B, E) Comparison of species and OTU richness between DNA metabarcoding and eDNA metabarcoding, with statistical significance tested using Kruskal–Wallis tests (**** $p < 0.0001$). (C, F) Species and OTU richness across habitats (degraded forest, grassland, limestone forest), with pairwise comparisons indicated by significance levels (** $p < 0.01$; ns = not significant).

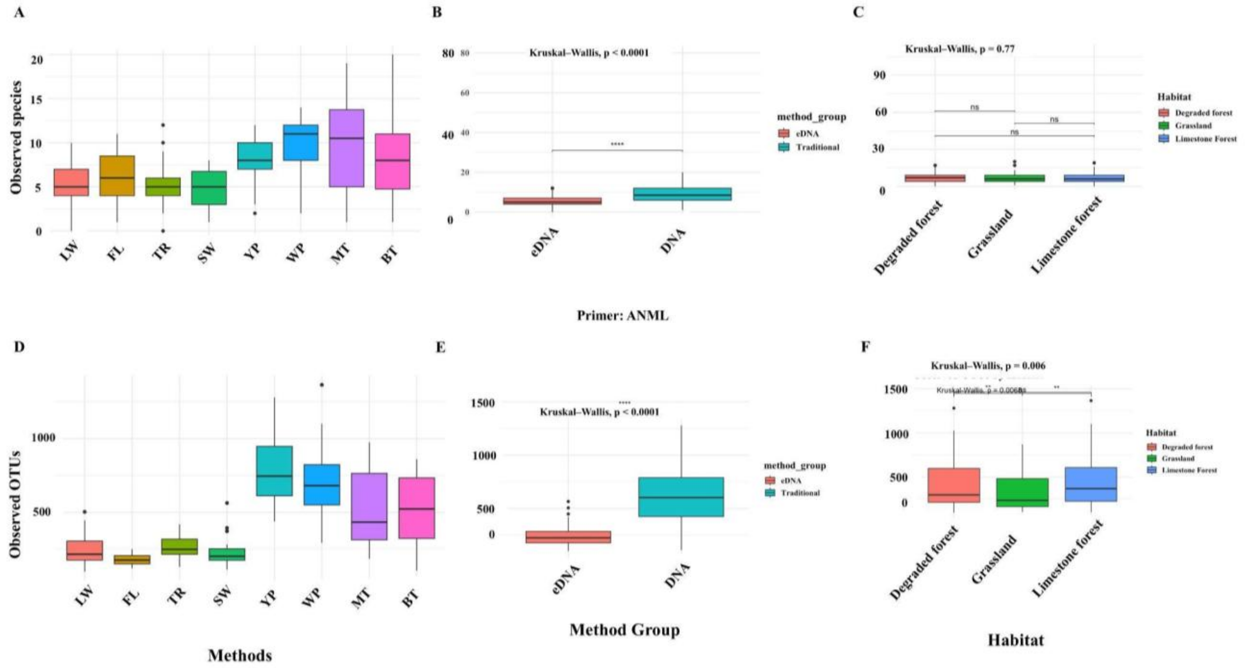


**Figure S6.** Alpha diversity of species (A–C) and OTUs (D–F) detected using the ANML primer across different methods, method groups, and habitats. (A, D) Boxplots showing the number of observed species and OTUs across individual sampling methods (MT: malaise traps, BT: beating sheets, WP: White Pan, YP: Yellow Pan, FL: flower, LW: leaf wash, TR: trunk rolling, and SW: spider webs). (B, E) Comparison of species and OTU richness between DNA metabarcoding and eDNA metabarcoding, with statistical significance tested using Kruskal–Wallis tests (**** $p < 0.0001$). (C, F) Species and OTU richness across habitats (degraded forest, grassland, limestone forest), with pairwise comparisons indicated by significance levels (** $p < 0.006$; ns = not significant).

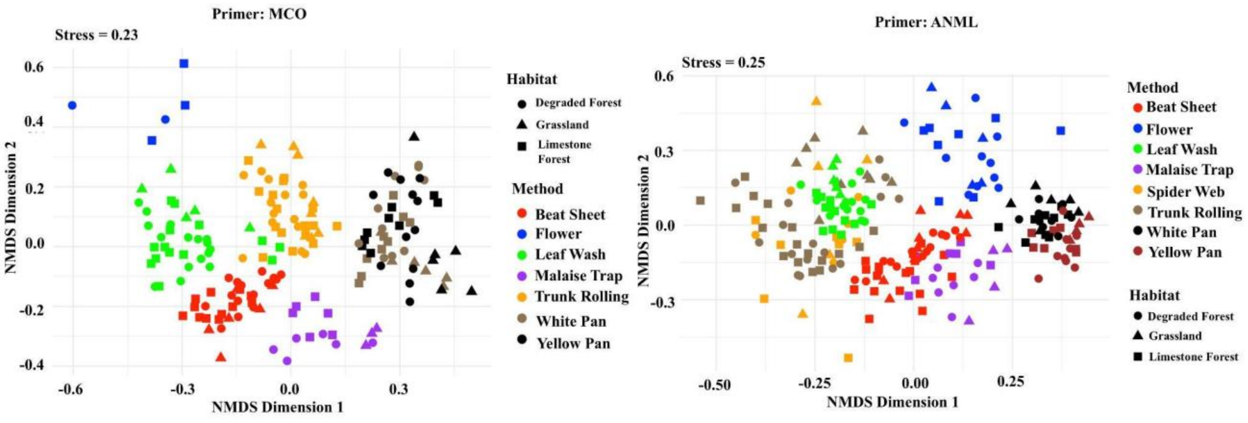


**Figure S7.** Beta diversity analysis based on non-metric multidimensional scaling (NMDS) ordination of community composition detected using the MCO primer (left) and ANML primer (right). Each point represents a sample, with colors indicating different sampling methods (Beat Sheet, Flower, Leaf Wash, Malaise Trap, Spider Web, Trunk Rolling, White Pan, Yellow Pan), and shapes representing habitats (degraded forest = circles, grassland = triangles, limestone forest

= squares). Stress values (0.23 for MCO; 0.25 for ANML) indicate acceptable representation of community dissimilarity in two dimensions. Clustering patterns illustrate differences in arthropod community composition across methods and habitats between the two primer sets.

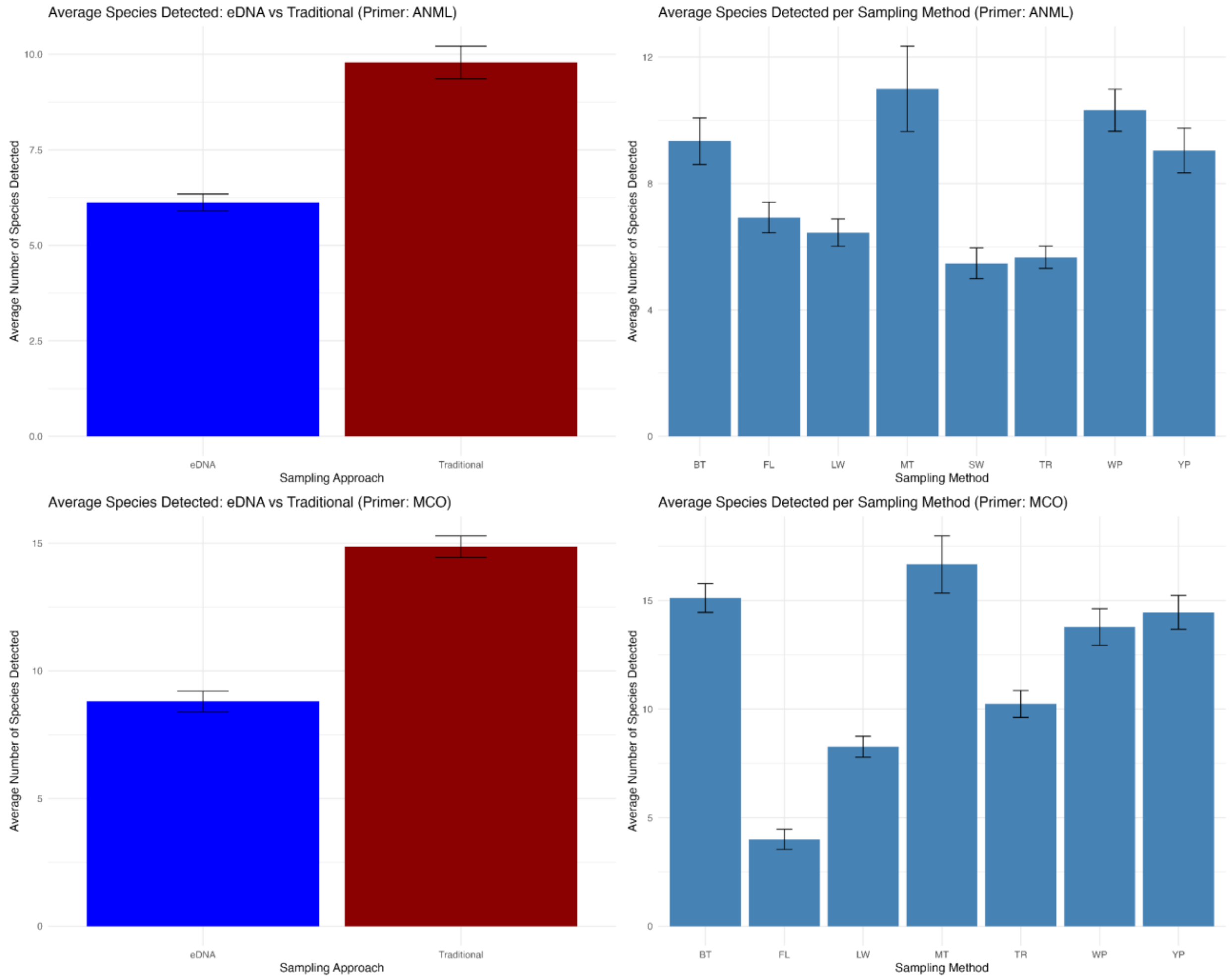


Figure S6. Average number of species per method and average number of species per method group. In the ANML primer, among all methods, the malaise trap detected the highest number of species per sample (11±1.3), followed by white pan (10.3±0.7), beating sheet (9.3±0.7), yellow pan (9.0±0.7), the flower (6.9±0.5), the leaf wash (6.4±0.4), trunk rolling (5.7±0.3), and spider web (5.5±0.5). On average, the bulk DNA metabarcoding method detected the highest number of species per sample (9.8 ± 0.4), compared to eDNA metabarcoding method (6.1 ± 0.2). Similar trends were noticed in MCO primers. The malaise trap detected the highest number of species per sample (16.7±1.3), followed by the beating sheet (15.1±0.7), yellow pan (14.5±0.8), white pan (13.8±0.8), trunk rolling (10.2±0.6), leaf wash (8.3±0.5), and flower (4±0.5). Similar to the ANML primer, on average, the bulk DNA metabarcoding method detected the highest number of species per sample (14.9±0.4), compared to the eDNA metabarcoding method (8.8±0.2) (Figure S6). Moreover, in comparison to primer-based detection, MCO detected more species per sample than the ANML primer.

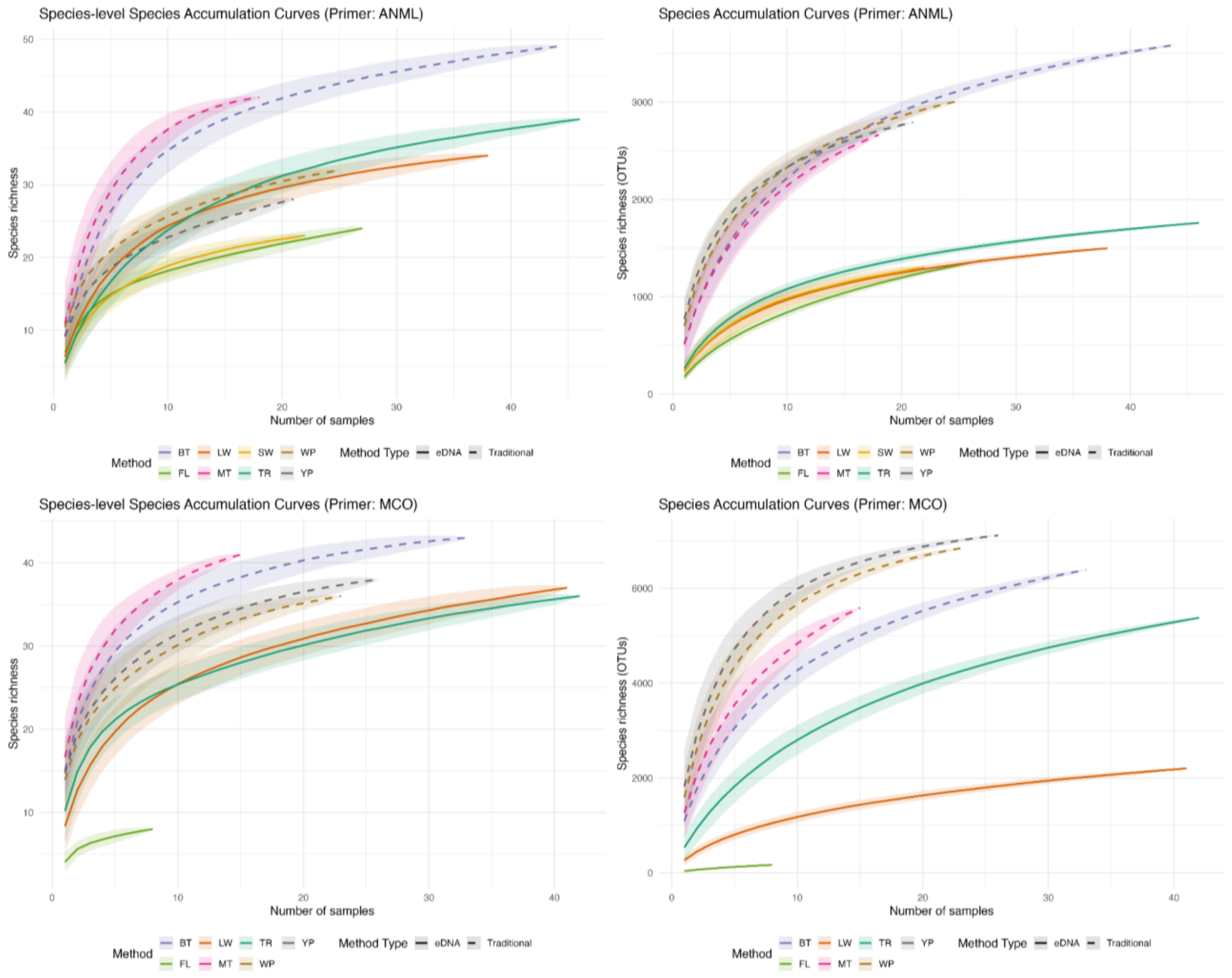


Figure S7. Species accumulation curves for arthropod communities assessed using different sampling methods and primers across Guam (top panels: ANML primer; bottom panels: MCO primer). Curves depict species richness as a function of the number of samples accumulated, with shaded ribbons representing standard deviations across permutations (999 permutations, random method). Solid lines indicate eDNA-based methods (TR, FL, LW, SW), while dashed lines represent traditional sampling methods (BT, MT, WP, YP). Left panels show species-level accumulation curves based on taxonomically assigned species abundances, highlighting biologically meaningful diversity estimates. Right panels display OTU-level accumulation curves, reflecting raw operational taxonomic unit diversity without taxonomic assignment bias.
